# POPEYE directly regulates bHLH Ib genes and its own expression

**DOI:** 10.1101/2022.03.09.482262

**Authors:** Meng Na Pu, Gang Liang

## Abstract

Iron (Fe) is an essential trace element for plants. When suffering from Fe deficiency, plants modulate the expression of Fe deficiency responsive genes. POPEYE (PYE) is a key bHLH transcription factor involved in Fe homeostasis. However, the molecular mechanism of PYE regulating the Fe deficiency response remains elusive. We found that the over-expression of *PYE* attenuates the expression of Fe deficiency responsive genes. PYE directly represses the transcription of bHLH Ib genes (*bHLH38, bHLH39, bHLH100*, and *bHLH101*) by associating with their promoters. Although PYE contains an <u>E</u>thylene response factor-associated <u>A</u>mphiphilic <u>R</u>epression (EAR) motif, it does not interact with the transcriptional corepressors TOPLESS/TOPLESS-RELATED (TPL/TPRs). Subcellular localization analysis indicated that PYE localizes in both the cytoplasm and nucleus. PYE contains a <u>N</u>uclear <u>E</u>xport <u>S</u>ignal (NES) which is required for the cytoplasmic localization of PYE. The mutation of NES amplifies the repression function of PYE, resulting in downregulation of Fe deficiency responsive genes. Co-expression assays indicated that bHLH IVc members (bHLH104, bHLH105/ILR3, and bHLH115) facilitate the nuclear accumulation of PYE. Conversely, PYE indirectly represses transcription activation ability of bHLH IVc. Additionally, PYE directly negatively regulates its own transcription. This study provides insights into the complicated Fe deficiency response signaling pathway and enhances the understanding of PYE functions.

**Short summary:** PYE is a negative regulator of Fe homeostasis; however, it was still unclear how PYE integrates the Fe deficiency response signaling. Our study shows that conditional nuclear localization of PYE is crucial for Fe homeostasis. PYE not only negatively regulates FIT-dependent Fe uptake genes by directly targeting bHLH Ib genes, but also negatively regulates its own expression.

## Introduction

Fe is an essential trace element for all biological organisms. It participates in not only the intracellular redox reactions, but also the electron transmission. In plants, Fe participates in cellular respiration, photosynthesis and catalytic reactions of metal proteins (Kobayashi and Nishizawa, 2014). On the other hand, due to the high redox potential of Fe^3+^/Fe^2+^, the free Fe ions in the cell are prone to Fenton reaction, which activates the reduced oxygen and produces harmful superoxide compounds causing damage to cells (Sheftel et al., 2012; Dixon and Stockwell, 2014). Therefore, plants must maintain Fe homeostasis through a rigorous set of regulatory mechanisms.

Although Fe is abundant in the earth’s crust, it mostly exists in the form of Fe^3+^, and its solubility is extremely low in high pH value and calcareous soil, which seriously affects its utilization efficiency. As soil salinization increases, Fe deficiency in plants becomes prevalent. In the long-term evolution process, plants have formed a set of sophisticate molecular mechanism for Fe absorption (Römheld and Marschner, 1986; Kobayashi and Nishizawa, 2012; Grillet and Schmidt, 2019). Dicotyledonous plants and non-graminaceous monocotyledonous plants absorb Fe by a reduction-based strategy which consists of three components, H^+^-ATPase, Fe^3+^ reduction enzyme and Fe^2+^ transporter. In *Arabidopsis*, the H^+^-ATPase AHA2 secretes H^+^ to reduce soil pH and increases Fe solubility in the rhizosphere (Santi and Schmidt, 2009), the Fe^3+^ reductase FRO2 (FERRIC REDUCTION OXIDASE 2) converts Fe^3+^ to Fe^2+^ (Robinson et al., 1999), and the Fe^2+^ transporter IRT1 (IRON-REGULATED TRANSPORTER 1) transports Fe^2+^ to root cells (Henriques et al., 2002; Varotto et al., 2002; Vert et al., 2002). Graminaceous plants utilize a chelation-based strategy in which low molecular weight mugineic acids are secreted into the rhizosphere to bind Fe^3+^ and then the chelation complex is transported to plant roots (Roberts et al., 2004).

When exposed to Fe deficiency conditions, plants activate their Fe uptake systems. A series of transcription factors have been characterized to regulate the Fe deficiency response (Gao and Dubos, 2021; Riaz and Guerinot, 2021). FER-LIKE IRON DEFICIENCY-INDUCED TRANSCRIPTION FACTOR (FIT) is the master regulator of *Arabidopsis* strategy I associated genes, such as *IRON-REGULATED TRANSPORTER 1* (*IRT1*) and *FERRIC REDUCTION OXIDASE 2* (*FRO2*), since their expression is compromised in the *fit* loss-of-function mutant which cannot survive without extra Fe supplementation (Colangelo and Guerinot, 2004; Jakoby et al., 2004; Yuan et al., 2005; Schwarz and Bauer, 2020). However, FIT alone is not sufficient to induce the expression of *IRT1* and *FRO2*. Four bHLH Ib subgroup members (bHLH38, bHLH39, bHLH100, and bHLH101) interact with FIT, and the co-overexpression of FIT and bHLH Ib constitutively activates the expression of *IRT1* and *FRO2* (Yuan et al., 2008; Wang et al., 2013). A recent study revealed that FIT has the transactivation activity and bHLH Ib members have the DNA binding ability, and they complement each other to from a functional transcription complex which initiates the expression of Fe-uptake genes (Cai et al., 2022). Both *FIT* and bHLH Ib genes are inducible in response to Fe deficiency. Four bHLH IVc proteins, bHLH34/IRON DEFICIENCY TOLERANT1 (IDT1), bHLH104, bHLH105/IAA-LEUCINE RESISTANT3 (ILR3), and bHLH115, directly bind to the promoters of bHLH Ib genes and promote their expression (Zhang et al., 2015; Li et al., 2016; Liang et al., 2017). As positive regulators of the Fe deficiency response, bHLH IVc members are not stimulated at the transcription level by Fe deficiency (Zhang et al., 2015; Li et al., 2016; Liang et al., 2017). In fact, bHLH IVc proteins are regulated at the post translation level as BRUTUS (BTS), a candidate of Fe sensor, promotes their degradation (Selote et al., 2015; Xing et al., 2020; Li et al., 2021a). Despite of the increase of BTS protein stability under Fe deficient conditions, a class of small peptides, IRONMANs, inhibit the interactions between bHLH IVc (bHLH105/bHLH115) and BTS, resulting in the elevation of bHLH IVc proteins (Li et al., 2021a).

One member of bHLH IVb subgroup, bHLH121/UPSTREAM REGULATOR of IRT1 (URI), also directly associates with the promoters of bHLH Ib genes and forms heterodimers with bHLH IVc members to activate bHLH Ib genes (Kim et al., 2019; Gao et al., 2020; Lei et al., 2020). The induction of *FIT* is blocked in the *bhlh121*/*uri* loss-of-function mutants (Kim et al., 2019; Gao et al., 2020; Lei et al., 2020), and the *FIT* promoter is also bond by bHLH121 (Lei et al., 2020). In addition to bHLH121, bHLH IVb subgroup also contains bHLH11 and POPEYE (PYE/bHLH47). bHLH11 is a negative regulator containing an EAR motif which can recruit the transcriptional corepressors TOPLESS/TOPLESS-RELATED (TPL/TPRs) (Tanabe et al., 2019; Li et al., 2021b). Moreover, bHLH11 interacts with bHLH IVc proteins and interferes with their transactivation to bHLH Ib genes (Li et al., 2021b). In contrast, PYE negatively regulates the expression of *NICOTIANAMINE SYNTHASE 4* (*NAS4*), *FRO3*, and *ZINC-INDUCED FACILITATOR1* (*ZIF1*) by directly binding to their promoters (Long et al., 2010). Although PYE is a negative regulator of Fe deficiency inducible genes, its loss-of-function mutant displays the enhanced sensitivity to Fe deficiency. Similar to other bHLH Ib members, PYE interacts with three bHLH IVc members (bHLH104, bHLH105, and bHLH115) (Long et al., 2010; Selote et al., 2015; Tissot et al., 2019); however, the biological significance of their protein interactions is still unclear. In the present study, we show that the cytoplasmic localization of PYE is required for the maintenance of Fe homeostasis. PYE represses the expression of bHLH Ib genes in both direct and indirect manners. Moreover, the transcription of *PYE* is also under the control of PYE itself.

## Results

### Overexpression of *PYE* suppresses both FIT-dependent and FIT-independent Fe deficiency response

To further investigate the functions of PYE in the Fe deficiency response, we constructed *PYE* overexpressing plants (*PYEox*), in which HA tagged *PYE* was driven by the CaMV 35S promoter (Supplemental Figure S1). Under Fe sufficient conditions, no visible difference was observed between the *PYEox* plants and the wild type. In contrast, under Fe deficient conditions, the *PYEox* plants displayed sensitivity to Fe deficiency, such as short roots (Figure 1A). We then analyzed the Fe reductase activity (Yi and Guerinot, 1996). The results indicated that the Fe reductase activity is lower in the *PYEox* plants than in the wild type (Figure 1B). These data suggest that *PYE* overexpression disrupts the Fe deficiency response.

**Figure 1.**
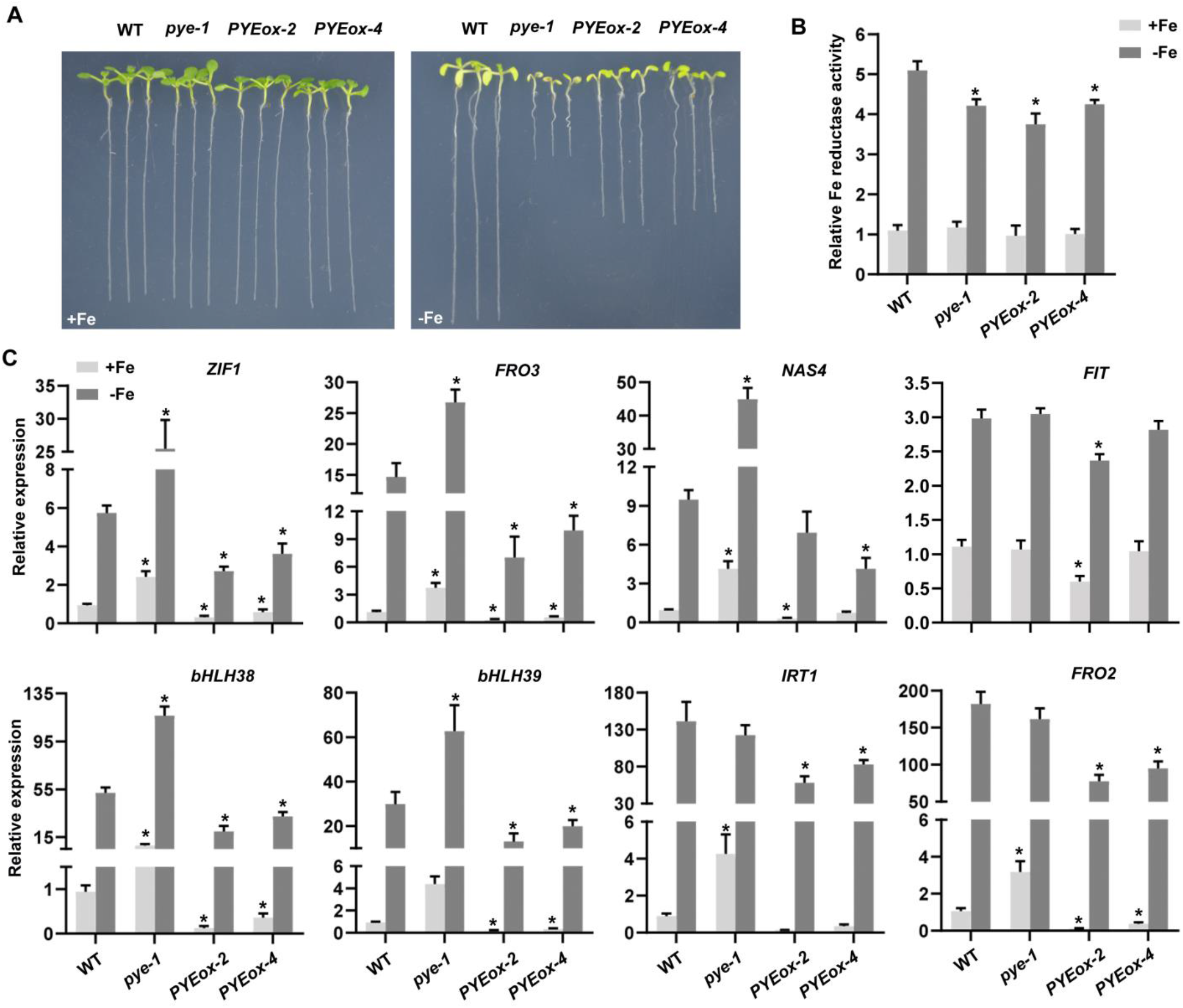
Characterization of *PYE* overexpressing plants. (A) Phenotypes of *PYEox* plants. One-week-old seedlings grown on Fe sufficient (+Fe) or Fe deficient (-Fe) medium. (B) Fe reductase activity. Four-day-old plants grown on +Fe were transferred to +Fe or -Fe medium for three days. The ferrozine assay was performed, in triplicate, on 10 pooled plant roots. The data represent means ± SD. The asterisks indicate that the values are significantly different from the corresponding wild type (WT) value by Student’s t Test (*P* < 0.05). (C) Expression of Fe deficiency responsive genes. Four-day-old plants grown on +Fe were transferred to +Fe or -Fe medium for three days, and root samples were used for RT-qPCR. The data represent means ± SD. The asterisks indicate that the values are significantly different from the corresponding wild type value by Student’s t Test (*P* < 0.05).

Next, we wondered whether the transcription of Fe deficiency response genes is also affected in the *PYEox* plants. qRT-PCR was used to examine the expression of genes involved in the Fe deficiency response, including *ZIF1, FRO3, NAS4, FIT, bHLH38, bHLH39, IRT1*, and *FRO2*, (Figure 1C). In agreement with the previous study (Long et al., 2010), the expression of *FIT, IRT1*, and *FRO2* was not significantly changed in the *pye-1*. In contrast, all these genes were significantly down-regulated in the *PYE*ox plants irrespective of Fe status. These results suggest that the overexpression of PYE downregulates not only the FIT-independent genes, but also the FIT-dependent genes.

### PYE directly regulates bHLH Ib genes

*PYE* overexpression significantly represses the expression of bHLH Ib genes whereas the loss-of-function of *PYE* causes the opposite result (Figure 1C), indicating that PYE negatively regulates bHLH Ib genes. Four bHLH IVc proteins are the positive regulators of bHLH Ib genes, and three of them physically interact with PYE (Long et al., 2010). It is possible that PYE inhibits the positive regulation of bHLH IVc proteins to bHLH Ib genes. However, it cannot exclude the possibility that PYE alone directly regulates bHLH Ib genes. To verify this hypothesis, we carried out transient expression assays (Figure 2A). The promoters of *bHLH38* and *bHLH39* were fused with a nucleus localized GFP (nGFP) as the reporters, *Pro*_*bHLH38*_*:nGFP* and *Pro*_*bHLH39*_*:nGFP*. The mCherry tag was fused with PYE and driven by the CaMV 35S promoter as an effector, *Pro*_*35S*_*:mCherry-PYE. Pro*_*35S*_*:mCherry- bHLH105* and *Pro*_*35S*_*:mCherry* were used as the positive and negative controls, respectively. Each reporter was co-expressed with each of the reporters respectively. Compared with mCherry, mCherry-PYE significantly inhibited the expression of *GFP* whereas mCherry-bHLH105 considerably promoted (Figure 2B). These results suggest that PYE alone is sufficient to represses bHLH Ib genes.

**Figure 2.**
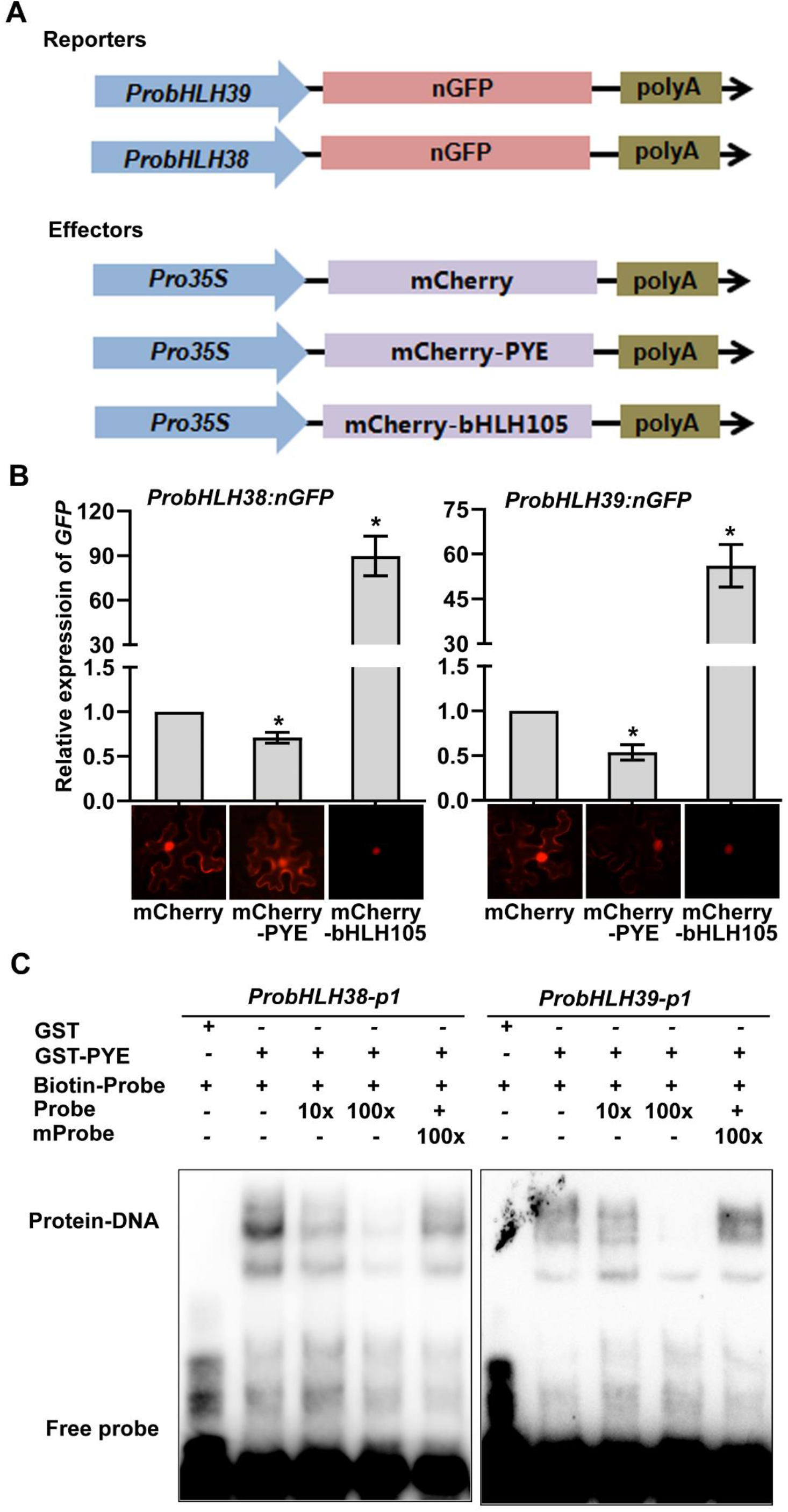
PYE directly regulates bHLH Ib genes. (A) Schematic representation of the constructs used for transient expression assays. The promoters of *bHLH38* and *bHLH3*9 were fused with a nucleus localized GFP (nGFP) as the reporters. The mCherry tag was fused with *PYE* and driven by the CaMV 35S promoter as an effector. *Pro*_*35S*_*:mCherry-bHLH105* and *Pro*_*35S*_*:mCherry* were used as the positive and negative controls, respectively. (B) *GFP* transcript abundance. Representative images for each mCherry tagged protein are shown. The abundance of *GFP* was normalized to that of *NPTII*. The value with the empty vector as an effector was set to 1. Each bar represents the mean ± SD of three independent experiments. The asterisks indicate that the values are significantly different from the corresponding wild type value by Student’s t Test (*P* < 0.05). (C) EMSA showing that PYE directly binds to the *bHLH38* and *bHLH39* promoters. Either GST-PYE or GST was incubated with the biotin-labeled probes. Biotin-Probe, biotin-labeled probe; Probe, unlabeled probe; mProbe, unlabeled probe with mutated E-box. Biotin probe incubated with GST served as the negative control.

It has been confirmed that four bHLH IVc proteins and bHLH121/URI directly bind to the promoters of bHLH Ib genes (Zhang et al., 2015; Li et al., 2016; Liang et al., 2017; Kim et al., 2019; Gao et al., 2020; Lei et al., 2020). Since both bHLH121 and PYE belong to the bHLH IVb subgroup (Heim et al., 2003), we speculated that PYE also directly regulates the transcription of bHLH Ib genes. To this aim, we conducted EMSAs (Figure 2C). The promoters of *bHLH38* and *bHLH39* were used as the probes. The GST tagged PYE recombinant protein was expressed and purified in *Escherichia coli*. The biotin labeled probe was incubated with GST-PYE and the shifted DNA-protein complexes was detected. When excessive probe without biotin was added, the number of DNA-protein complexes decreased dramatically. In contrast, the addition of mutant probes did not cause the reduction of DNA-protein complexes. As the negative control, GST protein alone could not bind to the promoters of *bHLH38* and *bHLH39*. Taken together, these data suggest that PYE directly represses the transcription of *bHLH38* and *bHLH39* by association with their promoters.

### PYE has a non-functional EAR motif

Having confirmed that PYE negatively affects the expression of Fe homeostasis associated genes, we wondered how PYE exerts its negative regulation function. It has been hypothesized that PYE might recruit the co-transcription repressors, TPL/TPRs (Szemenyei et al., 2008; Pauwels et al., 2010; Causier et al., 2012), since it has an EAR motif (DLNxxP) in the C-terminal region (Figure 3A). To confirm this hypothesis, we employed yeast-two-hybrid assays to test their protein interactions (Figure 3B). The N-terminal regions of TPL/TPRs fused with the BD were used as the baits, and PYE with the AD as the prey. bHLH11 with the AD was a positive control (Li et al., 2021b). Yeast growth indicated that PYE cannot interact with all TPL/TPRs (Figure 3B).

**Figure 3.**
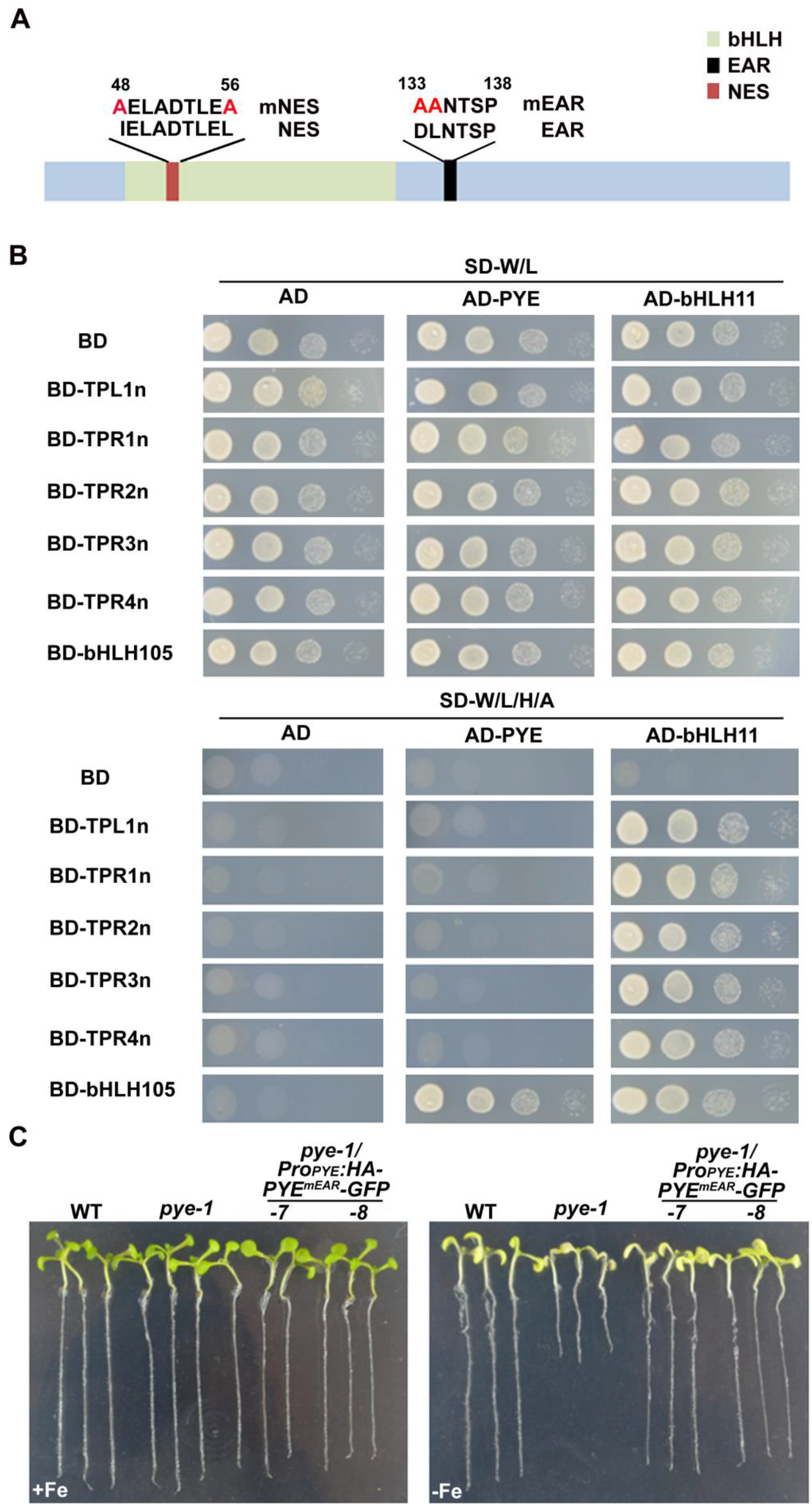
The repression function of PYE is independent of the EAR motif. (A) Schematic diagram of the various mutated versions of PYE. The mutated amino acids are indicated in red. mNES, the mutated NES. mEAR, the mutated EAR. (B) Yeast two-hybrid assays. Yeast co-transformed with different BD and AD plasmid combinations was spotted in parallel in a 10-fold dilution series. Growth on selective plates lacking Leu/Trp (SD–W/L) or Trp/Leu/His/Ade (SD–W/L/H/A). bHLH11 was used as the positive control for TPL/TPRs, and bHLH105 for PYE. (C) Complementation of the *pye-1* mutant by *PYE*^*mEAR*^. *PYE*^*mEAR*^ was fused with HA and GFP and driven by the *PYE* promoter. One-week-old seedlings grown on +Fe or -Fe medium.

To further investigate whether the EAR motif is responsible for the negative regulation of PYE, we generated a mutated version of PYE (PYE^mEAR^, a PYE version containing a mutated EAR motif). *PYE*^*mEAR*^ was fused with HA and GFP and driven by the *PYE* promoter. The *Pro*_*PYE*_*:HA-PYE*^*mEAR*^*-GFP* was transformed into the *pye-1* mutant plants. No matter under Fe sufficient conditions or under Fe deficient conditions, the *pye-1* mutant was completely rescued (Figure 3C).

To verify whether the EAR motif is required for the repression function of PYE, we constructed the *PYE*^*mEAR*^ overexpressing plants (*PYE*^*mEAR*^*ox*), in which HA tagged *PYE*^*mEAR*^ was driven by the CaMV 35S promoter (Supplemental Figure S2A, B). Under Fe deficient conditions, the Fe deficiency symptoms in the *PYE*^*mEAR*^*ox* plants were as severe as those in the *PYEox* plants (Supplemental Figure S2C). We also added an LxLxL type of EAR motif to the C-end of PYE and generated the *PYE*^*EAR*^*ox* plants (Supplemental Figure S2A, D). There were no visible differences between the *PYE*^*EAR*^*ox* and *PYEox* plants (Supplemental Figure S2C). Taken together, our data suggest that PYE cannot recruit the TPL/TPRs corepressors, and its EAR motif is not required for its repression function.

### PYE contains a Nuclear Export Signal (NES) which is required for its functions

We noticed that the PYE-mCherry proteins exist in both the cytoplasm and nucleus (Figure 2B). To validate this observation, we generated the PYE-GFP construct and performed transient expression assays in tobacco leaves and *Arabidopsis* protoplast cells. The results indicated that PYE-GFP fusion proteins are also localized in the cytoplasm and nucleus (Figure 4A; Supplemental Figure S3A). We also generated and examined the *Pro*_*PYE*_*:HA-PYE-GFP*/*pye-1* plants, finding that PYE-GFP signal is visible in both the cytoplasm and nucleus under Fe sufficient conditions, and it tends to accumulate in the nucleus under Fe deficient conditions (Figure 4B).

**Figure 4.**
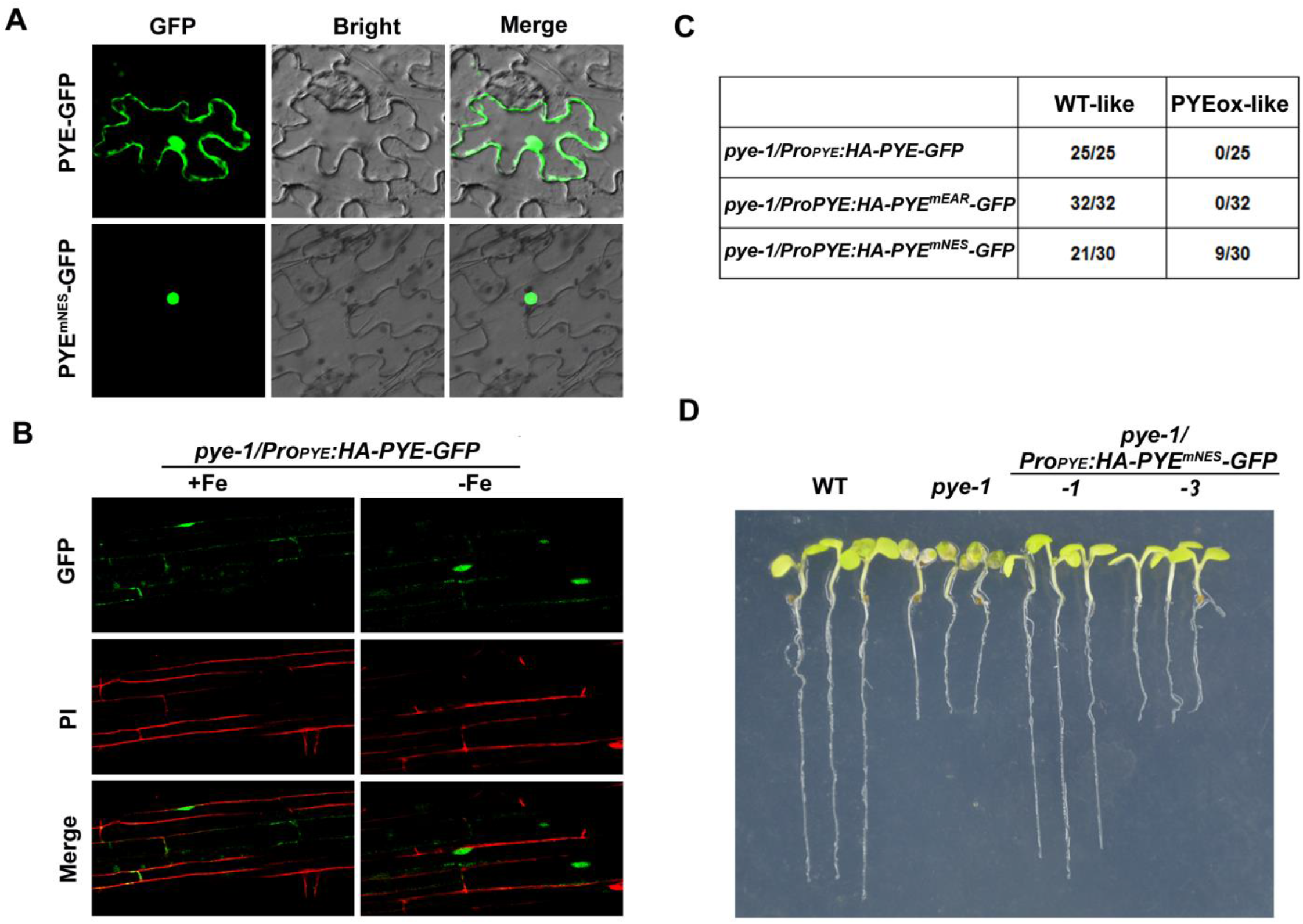
The NES is required for PYE functions. (A) Subcellular localization of PYE. PYE-GFP and PYE^mNES^-GFP were expressed in tobacco cells. The GFP signal was visualized under a confocal microscope. (B) Subcellular localization of PYE in response to Fe deficiency. One-week-old *Pro*_*PYE*_*:HA-PYE-GFP/pye-1* seedlings grown on +Fe or -Fe medium. Roots were stained with propidium iodide (PI, in red). Root maturation regions are shown. (C) Statistical analysis of the various versions of PYE plants under the *pye-1* background. (D) *Pro*_*PYE*_*:HA- PYE*^*mNES*^ *-GFP/pye-1* plants. One-week-old seedlings grown on -Fe medium.

A nuclear export signal (NES) consists of regularly spaced hydrophobic residues with several kinds of consensus patterns, and facilitates protein nuclear export (Xu et al., 2015). We employed the NetNES tool (la Cour et al., 2004) to predict the potential NES site of PYE, finding that PYE contains a NES site in the N-terminus (Figure 3A). To clarify whether the NES has an effect on PYE subcellular localization, we generated the PYE^mNES^-GFP (a PYE version containing a mutated NES) construct. The transient expression assays in tobacco leaves indicated that PYE^mNES^-GFP is predominantly localized in the nucleus (Figure 4A). To investigate whether the change of subcellular location of PYE would affect its function, we performed complementation assays. *Pro*_*PYE*_*:PYE*^*mNES*^*-GFP* was introduced into the *pye-1* mutant plants (Supplemental Figure S3B). Unlike *Pro*_*PYE*_*:PYE-GFP* and *Pro*_*PYE*_*:PYE*^*mEAR*^*-GFP*, both of which completely rescued the *pye-1* mutant (Supplemental Figure S3C), 70% of *Pro*_*PYE*_*:PYE*^*mNES*^*-GFP* transformants (WT-like) developed as well as the wild type whereas 30% of them (PYEox-like) displayed the Fe deficiency phenotypes as observed in the *PYEox* plants (Figure 4C, D; Supplemental Figure S3D). We then examined the Fe deficiency responsive genes, finding that their expression is lower in the *Pro*_*PYE*_*:PYE*^*mNES*^*-GFP/pye-1* than in the wild type (Figure 5). These findings suggest that *Pro*_*PYE*_*:PYE*^*mNES*^*-GFP* reverses the upregulation of Fe deficiency responsive genes in *pye-1*, but displays a strong repression function.

**Figure 5.**
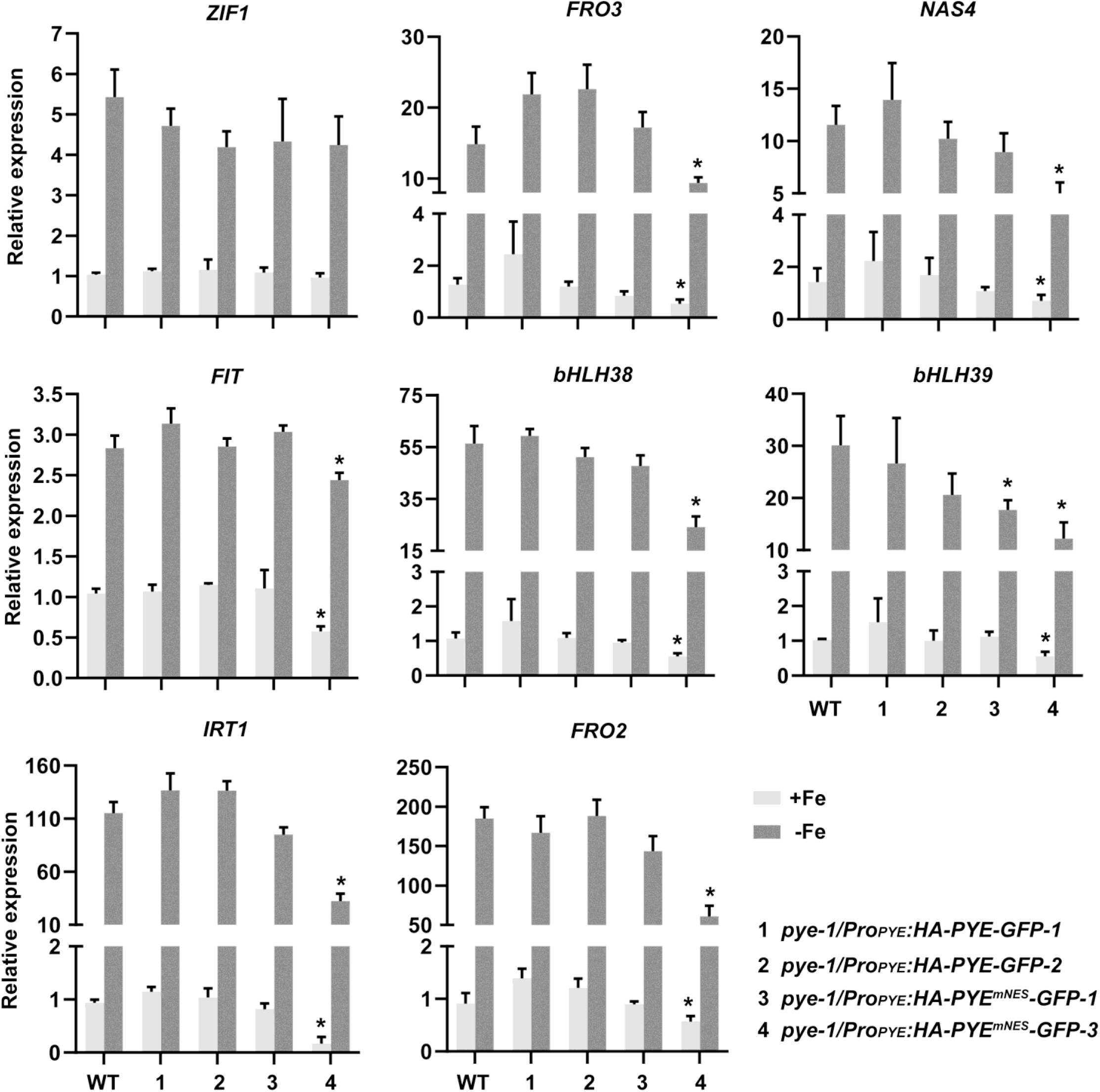
Expression of Fe deficiency responsive genes in *pye-1/Pro*_*PYE*_*:HA-PYE-GFP* and *pye-1/Pro*_*PYE*_*:HA-PYE*^*mNES*^*-GFP* plants. Plants were grown on +Fe medium for 4 days and then transferred to +Fe or -Fe medium for 3 days. RNA from root tissues was used for RT-qPCR. The data represent means ± SD. The asterisks indicate that the values are significantly different from the corresponding wild type value by Student’s t Test (*P* < 0.05).

### bHLH104, bHLH105, and bHLH115 promote the nuclear localization of PYE

Having confirmed that the NES is required for the cytoplasm localization of PYE, we then asked what facilitates the shuttle of PYE from cytoplasm to nucleus. It was reported that PYE can physically interact with three out of the four bHLH IVc proteins, bHLH104, bHLH105, and bHLH115, which are predominantly localized in the nucleus (Long et al., 2010; Selote et al., 2015; Lei et al., 2020). We wondered whether these bHLH IVc proteins contribute to the nuclear localization of PYE. The four GFP-tagged bHLH IVc proteins were respectively co-expressed with mCherry-PYE. We found that mCherry-PYE is mainly localized in the nucleus in the presence of bHLH104-GFP, bHLH105-GFP, or bHLH115-GFP (Figure 6). In contrast, the presence of GFP or bHLH34-GFP did not affect the subcellular localization of mCherry-PYE. Therefore, the PYE-interacting bHLH IVc proteins promote the nuclear localization of PYE.

**Figure 6.**
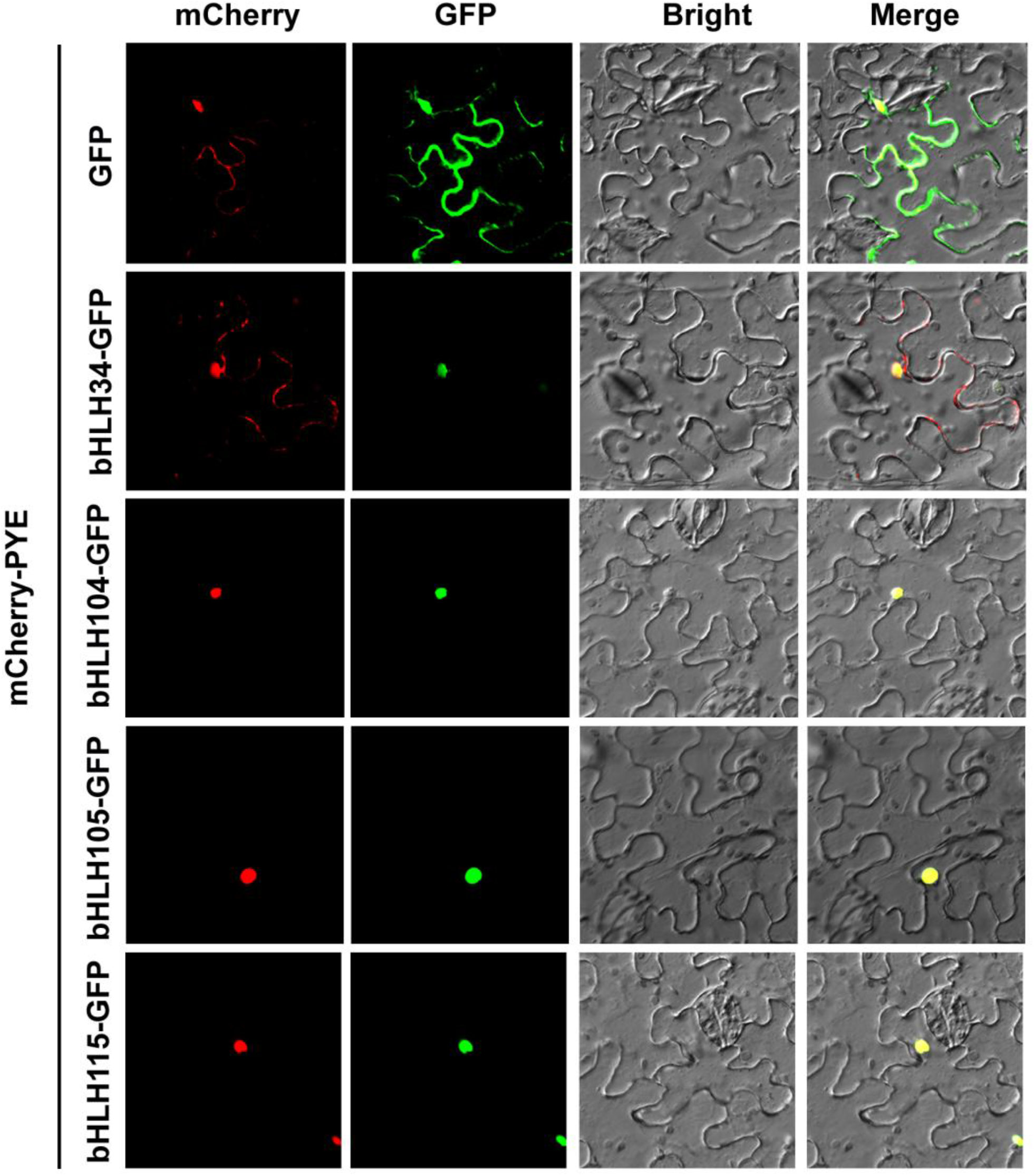
bHLH104, bHLH105 and bHLH115 promote the nuclear accumulation of PYE. The mCherry-PYE was co-transformed with GFP, bHLH34-GFP, bHLH104-GFP, bHLH105-GFP, or bHLH115-GFP into tobacco cells. GFP and mCherry signals were visualized under a confocal microscope.

### PYE represses the transcription activation ability of bHLH IVc

Given that PYE negatively regulates the expression of bHLH Ib genes, we further asked if PYE can antagonize the positive regulation function of bHLH IVc. Given the physically interaction between PYE and bHLH IVc (Long et al., 2010; Selote et al., 2015), we directly tested whether PYE inhibits the transcription activation of bHLH IVc by direct protein-protein interaction. We employed the GAL4-based effector-reporter system (Figure 7A). In the reporter, nGFP was driven by the minimal CaMV 35S promoter with five repeats of the GAL4 binding sequence. In the effectors, PYE, bHLH105, bHLH115 were fused with a nuclear localized mCherry (nmCherry) and the GAL4 DNA binding domain (BD) and driven by the 35S promoter. HA and HA-PYE were used as the secondary effectors. Consistent with the fact that bHLH IVc proteins are transcriptional activators and PYE is a repressor, the chimeric BD-nmCherry-bHLH105 and BD-nmCherry-bHLH115 activated the expression of *GFP* whereas BD-nmCherry-PYE inhibited. Compared with the control (HA), the co-expression of HA-PYE with BD-nmCherry-bHLH105 or BD-nmCherry-bHLH115 significantly reduced the expression of *GFP* (Figure 7B). Taken together, these data suggest that PYE antagonizes the transcriptional activation ability of bHLH IVc proteins.

**Figure 7.**
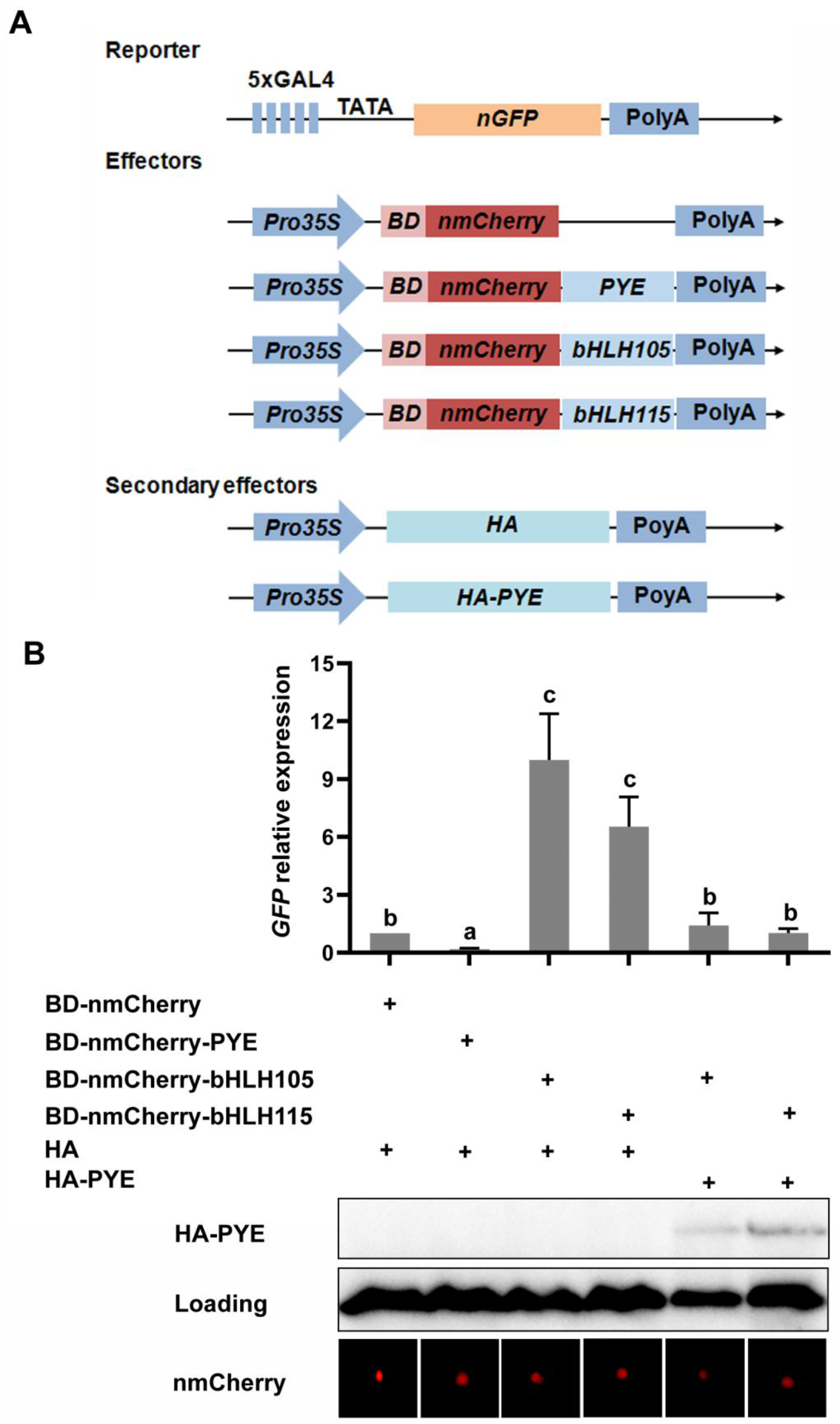
PYE represses the transcriptional activation ability of bHLH IVc. (A) The schematic diagram shows the constructs used in the transient expression assays. In the reporter, the minimal CaMV 35S promoter with five repeats of the GAL4 binding sequence drives the nuclear localized GFP (nGFP). In the effectors, the 35S promoter drives the fused genes in which the GAL4 DNA binding domain (BD), a target gene (PYE, bHLH105 or bHLH115) and the nuclear localized mCherry (nmCherry) were fused sequentially. HA and HA-PYE were used as the secondary effectors. (B) PYE represses transactivation of bHLH IVc. Protein levels of HA-PYE were detected by immunoblot. Representative images for each mCherry tagged protein were shown. The abundance of *GFP* was normalized to that of *NPTII*. The value with the empty vector as an effector was set to 1. Each bar represents the mean ± SD of three independent experiments. The different letters above each bar indicate statistically significant differences as determined by one-way ANOVA followed by Tukey’s multiple comparison test (P < 0.05).

### PYE directly regulates its own expression and physically interacts with itself

Both bHLH Ib genes and *PYE* are directly regulated by bHLH IVc members and bHLH121 (Zhang et al., 2014; Selote et al., 2015; Li et al., 2016; Liang et al., 2017; Kim et al., 2019; Gao et al., 2020; Lei et al., 2020). Having confirmed that PYE negatively regulates bHLH Ib genes, we wanted to know if PYE also negatively regulates its own expression. The *PYE* promoter was used to drive the GUS reporter, and then this construct (*Pro*_*PYE*_*:GUS*) was introduced into the wild type plants. GUS staining indicated that the *PYE* promoter is active in both the root and shoot (Figure 8A). The transgenic line *Pro*_*PYE*_*:GUS-2* was crossed with the *pye-1*, and the homozygous *pye-1*/*Pro*_*PYE*_*:GUS-2* line was identified. GUS staining analysis indicated that the *PYE* promoter activity is stronger in the *pye-1* than in the wild type (Figure 8B). To further quantify the activity of *PYE* promoter, we determined the expression levels of *GUS* gene. We found that the transcript levels of *GUS* are higher in the *pye-1* than in the wild type, irrespective of Fe status (Figure 8B), suggesting that *PYE* promoter activity is positively regulated by the PYE protein.

**Figure 8.**
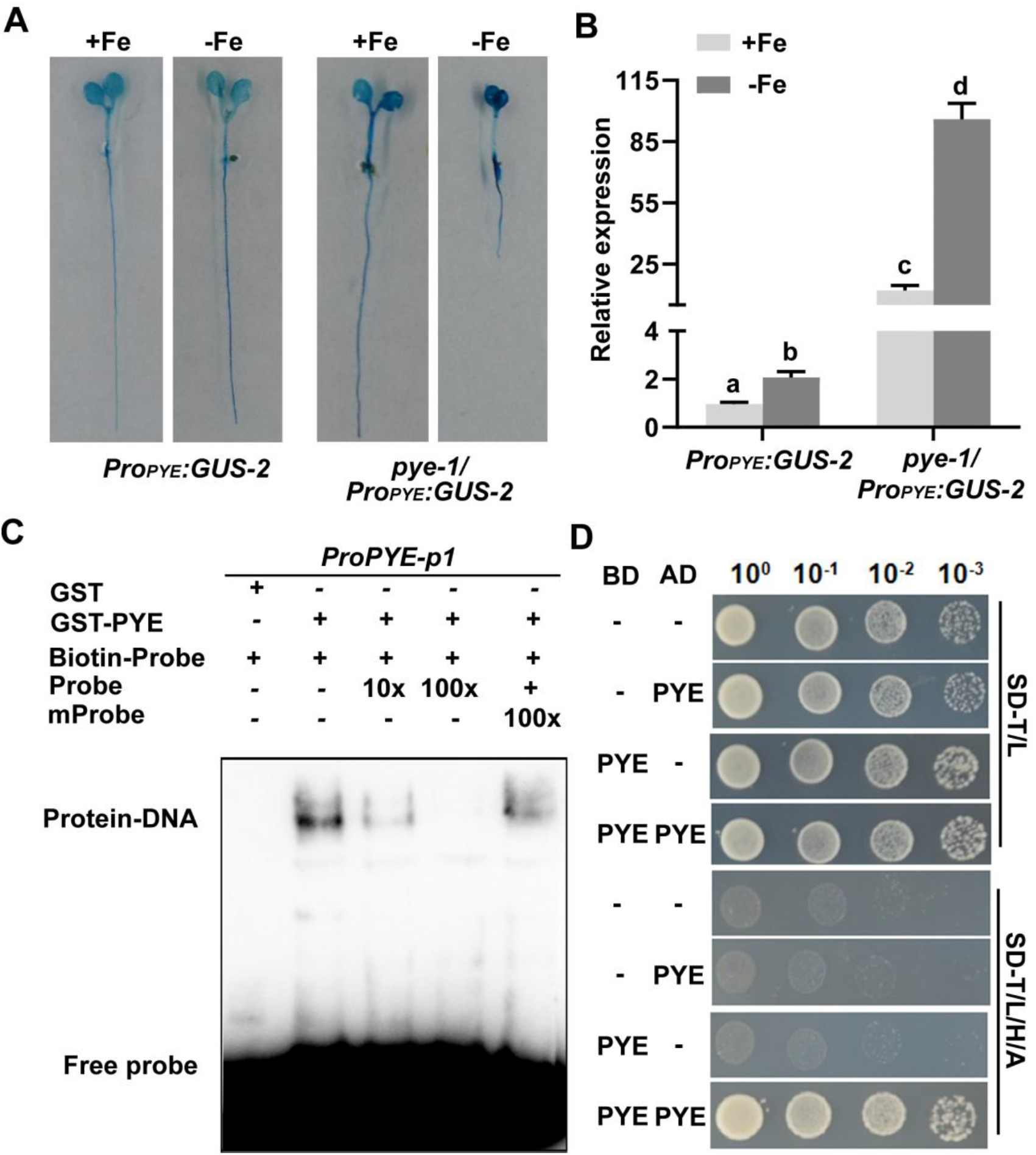
PYE directly regulates its own expression and physically interacts with itself. (A) GUS staining. One-week-old seedlings grown on +Fe or -Fe medium. Whole seedlings were subjected to GUS staining. (B) Expression of *GUS*. Four-day-old plants grown on +Fe were transferred to +Fe or -Fe medium for three days, and root samples were used for RT-qPCR. The data represent means ± SD. The different letters above each bar indicate statistically significant differences as determined by one-way ANOVA followed by Tukey’s multiple comparison test (P < 0.05). (C) EMSA showing that PYE directly binds the *PYE* promoter. Either GST-PYE or GST was incubated with the biotin-labeled probes. Biotin-Probe, biotin-labeled probe; Probe, unlabeled probe; mProbe, unlabeled probe with mutated E-box. Biotin probe incubated with GST served as the negative control. (D) Yeast two-hybrid assays. Yeast co-transformed with different BD and AD plasmid combinations was spotted in parallel in a 10-fold dilution series. Growth on selective plates lacking Leu/Trp (SD–W/L) or Trp/Leu/His/Ade (SD–W/L/H/A).

Next, we directly tested whether the PYE protein is able to bind to its own promoter. For this purpose, a fragment of *PYE* promoter with an E-box was subjected to EMSAs. The competition experiments showed that PYE specifically binds to the *PYE* probe (Figure 8C). Considering that PYE plays a strong inhibition role and directly binds to target promoters, we proposed that PYE can form homodimers. To confirm this hypothesis, we conducted yeast-two-hybrid assays. The results indicated that homodimerization occurs in PYE proteins (Figure 8D). Taken together, these results show that PYE is able to repress its own expression by directly binding to its promoter, and physically interacts with itself.

## Discussion

Fe deficiency is harmful to growth and development of plants. Plants can sense Fe deficiency conditions and modulate the expression of Fe deficiency responsive genes in order to maintain Fe homeostasis. Many transcription factors, especially the bHLH proteins, play pivotal regulatory roles in the Fe deficiency response signaling pathway (Gao and Dubos, 2021; Riaz and Guerinot, 2021). PYE was characterized as a direct negative regulator of the Fe homeostasis associated genes, *ZIF1, FRO3*, and *NAS4* (Long et al., 2010). Since loss-of-function of *PYE* does not change the expression of FIT target genes *IRT1* and *FRO2* under Fe deficient conditions, PYE was thought to regulate Fe homeostasis in an FIT-independent manner (Long et al., 2010; Riaz and Guerinot, 2021). Here, we provide evidence that PYE also directly represses bHLH Ib genes, resulting in the down-regulation of FIT-dependent Fe uptake genes. FIT is the master regulator of Fe uptake systems because it is crucial for the upregulation of Fe uptake genes, such as *IRT1* and *FRO2*, under Fe deficient conditions (Colangelo and Guerinot, 2004; Jakoby et al., 2004; Yuan et al., 2005; Schwarz and Bauer, 2020). Interestingly, we found that the overexpression of *PYE* causes the downregulation of both FIT-independent and FIT-dependent Fe deficiency responsive genes irrespective of Fe status. However, the expression of *FIT* is not or slightly affected in the *PYEox* plants (Figure 1C). We further confirmed that PYE directly represses the expression of bHLH Ib genes, *bHLH38* and *bHLH39*, by association with their promoters (Figure 2). It is well known that the bHLH Ib proteins interact with FIT to regulate the Fe uptake genes (Yuan et al., 2008; Wang et al, 2013). Recently, it was confirmed that bHLH Ib and FIT form a functional complex in which bHLH Ib accounts for binding to promoters of target genes and FIT for the transcription activation of target genes, namely, one of both cannot exert the regulatory function in the absence of the other one (Cai et al., 2022). We propose that the significant reduction of bHLH Ib genes is the reason that *IRT1* and *FRO2* are downregulated in the *PYEox* plants. Although the FIT-dependent Fe uptake genes are not affected in the *pye-1* mutant under Fe deficient conditions, their expression significantly increases under Fe sufficient conditions (Figure 1D). It is very likely that some unknown factors, which are increased in the *pye-1* mutant under Fe deficient conditions, facilitate the degradation of bHLH Ib or (and) FIT proteins, or interfere with their interaction. Further investigation is required to clarify why loss-of-function of *PYE* does not affect FIT-dependent Fe uptake genes under Fe deficient conditions.

Many negative regulators exert their repression function by recruiting transcriptional co-repressors. A recent study revealed that bHLH11, a homolog of PYE, functions as a repressor since its EAR motif can interact with the transcriptional co-repressors, TPL/TPRs. PYE is a negative regulator of Fe homeostasis. Due to the existence of an EAR motif in PYE, it was thought that PYE can recruit TPL/TPRs to inhibit its target gene transcription. However, our data negate this hypothesis. PYE, bHLH11 and bHLH121 belong to the bHLH IVb subgroup. It has been confirmed that bHLH11 contains an EAR motif which contributes to its negative function by recruiting TPL/TPRs (Li et al., 2021b). Although bHLH121 is the closet homolog of bHLH11, it is not a negative regulator (Kim et al., 2019; Gao et al., 2020; Lei et al., 2020). Thus, these three members have evolved different functions. Further exploring which factors contribute to the negative function of PYE will enhance our understanding of the Fe deficiency response signaling.

The balance of positive regulators and negative regulators is curial for the maintenance of Fe homeostasis. In contrast to the slight response of *FIT* transcription to Fe deficiency, the response to bHLH Ib genes is intensive. Due to the interdependence between bHLH Ib and FIT, modulating the expression of bHLH Ib genes is sufficient to control the FIT-dependent Fe uptake genes. As the major positive regulators of the Fe deficiency response, the bHLH IVc subgroup proteins, bHLH34, bHLH104, bHLH105, and bHLH115, directly activate bHLH Ib genes. bHLH121, one member of bHLH IVb subgroup, is also required for the upregulation of bHLH Ib genes, and bHLH121 directly binds to their promoters (Kim et al., 2019; Gao et al, 2020; Lei et al., 2020). By contrast, bHLH11 is a negative regulator of bHLH Ib genes since it inhibits the transcription activation activity of bHLH IVc towards bHLH Ib genes (Li et al., 2021b). Here, we suggest that PYE can directly represses bHLH Ib genes by binding to their promoters. Both PYE and bHLH IVc bind to the E-box *cis*-elements in the promoters of bHLH Ib genes (Figure 2C). Thus, it is likely that they compete with each other for binding to the promoters. On the other hand, PYE may indirectly repress bHLH Ib genes since it can repress the transcription activation ability of bHLH IVc (Figure 7B). Therefore, bHLH IVc and PYE antagonistically regulate the expression of bHLH Ib genes. The reciprocal antagonistic regulations of these transcription factors enable plants to modulate the expression levels of bHLH Ib genes, finally finetuning the Fe uptake. In addition to bHLH Ib genes, *PYE* gene also must be maintained at an appropriate level since the non-expression or over-expression of *PYE* causes the damage in the growth of plants. We show evidence that PYE directly represses its own expression. It is worthy to mention that bHLH IVc proteins directly activate the expression of *PYE* (Zhang et al., 2015; Liang et al., 2017). Therefore, the balance of bHLH IVc and PYE finally determines the expression levels of *PYE*.

The nuclear localization of transcription factors is required for them to exert regulatory function since they need to bind to target promoters in nuclei. It is noteworthy that all three bHLH IVb members localize in the cytoplasm and nucleus, and their interacting bHLH IVc proteins facilitate their retention in the nucleus (Figure 6; Lei et al., 2020; Li et al., 2021b). It remains unclear if the cytoplasmic localization of bHLH IVb proteins is related with their roles in the Fe deficiency response. Three bHLH IVc proteins promote the nuclear accumulation of PYE, and the nuclear PYE proteins increase under Fe deficient conditions. We show that the removal of the NES enables PYE^mNES^ to stay in the nucleus. When *PYE*^*mNES*^ was driven by the native promoter in *pye-1*, the expression of Fe deficiency responsive genes was repressed significantly, suggesting that constitutive nuclear localization of PYE is harmful to plants. Thus, conditional nuclear localization of PYE is crucial for Fe homeostasis. Under Fe deficient conditions, the expression of *PYE* is induced, which means that plants need its negative function to avoid the excessive increase of Fe homeostasis associated genes. Indeed, the loss of this “brake” enables plants not to survive Fe deficient conditions, as shown in the *pye-1* mutant (Long et al., 2010). In addition to bHLH IVb proteins, two bHLH Ib proteins, bHLH38 and bHLH39, are also preferentially expressed in the cytoplasm, and they exclusively stay in the nucleus in the presence of FIT (Trofimov et al., 2019; Cai et al., 2022). It is still unknown if their cytoplasm localization is required for their functions. In rice, the bHLH Ib protein, OsIRO2, also localizes in the cytoplasm, and its interacting partner OsFIT facilitates its accumulation in the nucleus (Wang et al., 2019; Liang et al., 2020). One member of bHLH IVb subgroup, OsIRO3, was reported to be localized in the nucleus (Zheng et al., 2010), and the subcelluar localization of the other members has not been reported. Further investigation is required to clarify whether the cytoplasm retention phenomenon of bHLH IVb proteins universally exists across different plant species and is crucial for Fe homeostasis.

## Materials and methods

### Plant materials and growth conditions

*Arabidopsis thaliana* ecotype Columbia-0 was used as the wild type in this study. *pye-1* was described previously (Long et al., 2010). Plants were grown in long photoperiods (16 h of light / 8 h of dark) or in short photoperiods (8 h of light / 16 h of dark) at 22°C. Surface-sterilized seeds were stratified at 4°C for 1 day before being planted on medium. Fe sufficient medium is one-half-strength Murashige and Skoog (MS) medium with 1% (w/v) Suc, 0.7% (w/v) agar A, and 0.1 mM FeEDTA. Fe-deficient medium is the same without FeEDTA.

### Plasmid construction

Standard molecular biology techniques were used for the cloning procedures. Genomic DNA from *Arabidopsis* was used as the template for amplification of the upstream regulatory promoter sequences of *PYE, bHLH38* and *bHLH39*. For *Pro*_*PYE*_:*HA*-*PYE-GFP, Pro*_*PYE*_:*HA*-*PYE*^*mEAR*^*-GFP* and *Pro*_*PYE*_:*HA*-*PYE*^*mNES*^*-GFP*, various versions of *PYE* sequences were inserted between the *PYE* promoter and poly(A) of the binary vector pOCA28. For overexpression of various versions of *PYE*, the HA tagged *PYE* (or *PYE*^*mEAR*^ or *PYE*^*mNES*^) was inserted between the CaMV 35S promoter and poly(A) of the binary vector pOCA30. Primers used for construction of these vectors are listed in the Supplemental Table S1. *Arabidopsis* transformation was conducted by the floral dip method (Clough and Bent, 1998). Transgenic plants were selected with the use of 50 μg mL^-1^ kanamycin.

### Fe reductase activity

Ferric chelate reductase assays were performed as described previously (Yi and Guerinot, 1996). Briefly, 10 intact plants for each genotype were pretreated for 30 min in plastic vessels with 4 ml of one-half-strength MS solution without micronutrients (pH 5.5) and then soaked into 4 ml of Fe (III) reduction assay solution [one-half-strength MS solution without micronutrients, 0.1 mM Fe (III)-EDTA, and 0.3 mM ferrozine, pH adjusted to 5 with KOH for 30 min in darkness. An identical assay solution containing no plants was used as a blank. The purple-colored Fe (II)-ferrozine complex was quantified at 562 nm.

### Gene expression analysis

One microgram of total RNA extracted using the Trizol reagent (Invitrogen) was used for oligo(dT)18-primed cDNA synthesis according to the reverse transcription protocol (TaKaRa). The resulting cDNA was subjected to relative quantitative PCR using the SYBR Premix Ex Taq kit (TaKaRa) on a Roche LightCycler 480 real-time PCR machine, according to the manufacturer’s instructions. For the quantification of each gene, at least three biological repeats were used. Gene copy number was normalized to that of *ACT2* and *PP2A*. Primers used for qRT-PCR were described previously (Li et al., 2021b).

### Subcellular localization

GFP and mCherry were cloned into pOCA30 to generate *Pro35S:GFP* and *Pro*_*35S*_*:mCherry*, respectively. PYE was fused with mCherry in *Pro*_*35S*_*:mCherry* to generate *Pro*_*35S*_*:mCherry-PYE. bHLH34, bHLH104, bHLH105*, and bHLH115 were cloned into *Pro*_*35S*_*:GFP* to generate *Pro*_*35S*_*:bHLH34-GFP, Pro*_*35S*_*:bHLH104-GFP, Pro*_*35S*_*:bHLH105-GFP*, and *Pro*_*35S*_*:bHLH115-GFP*, respectively (Lei et al., 2020). *Pro*_*35S*_*:mCherry-PYE* was co-expressed with various GFP containing vectors in *N. benthamiana* cells. Epidermal cells were observed under an Olympus confocal microscope. Excitation laser wavelengths of 488 nm and 563 nm were used for imaging GFP and mCherry signals, respectively.

### Transient expression assays in *Nicotiana benthamiana*

*Agrobacterium tumefaciens* strains EHA105 was used in the transient expression experiments in *N. benthamiana*. Agrobacterial cells were infiltrated into leaves of *N. benthamiana* by the infiltration buffer (0.2 mM acetosyringone, 10 mM MgCl_2_ and 10 mM MES, pH 5.6). In the transient expression assays, the final optical density at 600 nm value was 1. After infiltration 2 days in dark, GFP fluorescence was observed through a confocal laser scanning microscope, and leaf samples were harvested. *pGAL4* promoter and BD domain were described previously (Li et al., 2016). *pGAL4* promoter was fused with nGFP and cloned into the pOCA28 binary vector. *Pro35S:BD-mCherry, Pro35S:BD-mCherry-bHLH105, Pro35S:BD- mCherry-bHLH115*, and *Pro35S:HA-PYE* were constructed in the pOCA30 binary vector. For co-infiltration, different constructs were mixed prior to infiltration. Leaf infiltration was conducted in 3-week-old *N. benthamiana. NPTII* gene in the pOCA28 vector was used as the internal control. *GFP* transcript abundance was normalized to that of *NPTII*.

### Immunoblotting

For total protein isolation, samples were ground in liquid nitrogen and resuspended in RIPA buffer. Nuclear and cytosolic proteins were extracted as described previously (Saleh et al., 2008) with minor modification. In brief, *Arabidopsis* tissues were ground and resuspended in 2 ml of pre-chilled nuclei isolation buffer (0.25 M sucrose, 15 mM PIPES [pH 6.8], 5 mM MgCl_2_, 60 mM KCl, 15 mM NaCl, 1 mM CaCl_2_, 0.9% Triton X-100, and 13 protease inhibitor cocktail [Roche]). After homogenization, the slurry was filtered and centrifuged at 12 000 g for 20 min at 4°C. The proteins from the supernatant were extracted as cytosolic proteins, whereas the pellet resuspended in cold nuclei lysis buffer (50 mM HEPES [pH 7.5], 150 mM NaCl, 1 mM EDTA, 1% SDS, 0.1% Na deoxycholate, 1% Triton X-100, and 13 protease inhibitor cocktail [Roche]) was collected as nuclear proteins. Sample was loaded onto 12% SDS–PAGE gels and transferred to nitrocellulose membranes. The membrane was blocked with Tris buffer saline with Tween-20 (TBST) (10-mM Tris–Cl, 150-mM NaCl, and 0.05% [v/v] Tween-20, pH 8.0] containing 5% (w/v) nonfat milk (TBSTM) at room temperature for 60 min and incubated with primary antibody in Tris buffer saline with Tween-20 and milk (TBSTM) (overnight at 4°C). Membranes were washed with TBST (three times for 5 min each) and then incubated with the appropriate horseradish peroxidase conjugated secondary antibodies in TBSTM at room temperature for 1.5 h. After washing three times, bound antibodies were visualized with enhanced chemiluminescence (ECL) substrate.

### EMSA

EMSAs were conducted using the Chemiluminescent EMSA Kit (Beyotime, China) following the manufacturer’s protocol. The recombinant GST-PYE protein was purified from *E. coli*. The culture solution was incubated with 0.5 M isopropyl β-D-1-thiogalactopyranoside at 16°C for 16 h, and protein was extracted and purified by using the GST-tag Protein Purification Kit (Beyotime, China) following the manufacturer’s protocol. The DNA fragments of *bHLH38* and *PYE* promoters were synthesized and biotin was added to the 5’ terminus of the DNA. Unlabeled fragments of the same sequences or mutated sequences were used as competitors, and the GST protein alone was used as the negative control.

## Acknowledgements

We thank Germplasm Bank of Wild Species in Southwest China for confocal laser scanning microscopy.

## Finding

This work was supported by the National Natural Science Foundation of China (31770270).

## Author contributions

G.L. conceived the project. M.P. conducted all experiments. M.P. and G.L. wrote the manuscript, and approved the manuscript.

## Supplemental Information

**Supplemental Figure S1**. Relative expression levels of *PYE* in *PYEox* plants.

**Supplemental Figure S2**. The EAR of PYE is not required for its repression function.

**Supplemental Figure S3**. The NES is required for PYE functions.

**Supplemental Table S1**. Primers used in this study.

## References

Cai, Y., Yang, Y., Ping, H., Lu, C., Lei. R., Li, Y., Liang, G. (2022). Why FIT and bHLH Ib interdependently regulate Fe-uptake. bioRxiv 2022.02.12. 480172.

Causier, B., Ashworth, M., Guo, W., and Davies, B. (2012). The TOPLESS interactome: a framework for gene repression in Arabidopsis. Plant physiol. 158: 423–438.

Xu, D., Marquis, K., Pei, J., Fu, S. C., Cağatay, T., Grishin, N. V., and Chook, Y. M. (2015). LocNES: a computational tool for locating classical NESs in CRM1 cargo proteins. Bioinformatics 31: 1357–1365.

Dixon, S. J., and Stockwell, B. R. (2014). The role of Fe and reactive oxygen species in cell death. Nat Chem Biol. 10: 9–17.

Gao, F., Robe, K., Bettembourg, M., Navarro, N., Rofidal, V., Santoni, V., Gaymard, F., Vignols, F., Roschzttardtz, H., Izquierdo, E., and Dubos, C. (2020). The Transcription Factor bHLH121 Interacts with bHLH105 (ILR3) and Its Closest Homologs to Regulate Iron Homeostasis in Arabidopsis. Plant Cell 32: 508–524.

Gao, F., and Dubos, C. (2021). Transcriptional integration of plant responses to iron availability. J Exp Bot. 72: 2056–2070.

Heim, M. A., Jakoby, M., Werber, M., Martin, C., Weisshaar, B., and Bailey, P. C. (2003). The basic helix-loop-helix transcription factor family in plants: a genome-wide study of protein structure and functional diversity. Mol Biol Evol. 20: 735–747.

Henriques, R., Jásik, J., Klein, M., Martinoia, E., Feller, U., Schell, J., Pais, M. S., and Koncz, C. (2002). Knock-out of Arabidopsis metal transporter gene IRT1 results in iron deficiency accompanied by cell differentiation defects. Plant Mol Biol. 50: 587–597.

Kassebaum, N. J., Jasrasaria, R., Naghavi, M., Wulf, S. K., Johns, N., Lozano, R., Regan, M., Weatherall, D., Chou, D. P., Eisele, T. P., Flaxman, S. R., Pullan, R. L., Brooker, S. J., and Murray, C. J. (2014). A systematic analysis of global anemia burden from 1990 to 2010. Blood 123: 615–624.

Kobayashi T, Nishizawa N K. (2012). Fe Uptake, Translocation, and Regulation in Higher Plants. Ann.rev.plant Biol. 63: 131.

la Cour, T., Kiemer, L., Mølgaard, A., Gupta, R., Skriver, K., and Brunak, S. (2004). Analysis and prediction of leucine-rich nuclear export signals. Protein Eng Des Sel. 17: 527–536.

Liang, G., Zhang, H., Li, X., Ai, Q., and Yu, D. (2017). bHLH transcription factor bHLH115 regulates iron homeostasis in Arabidopsis thaliana. J Exp Bot. 68: 1743–1755.

Li, X., Zhang, H., Ai, Q., Liang, G., and Yu, D. (2016). Two bHLH Transcription Factors, bHLH34 and bHLH104, Regulate Iron Homeostasis in Arabidopsis thaliana. Plant Physiol. 170: 2478–2493.

Li Y, Lu CK, Li CY, Lei RH, Pu MN, Zhao JH, Peng F, Ping HQ, Wang D, Liang G. (2021a). IRON MAN interacts with BRUTUS to maintain iron homeostasis in Arabidopsis. Proc Natl Acad Sci USA. 118: e210906311.

Li, Y., Lei, R., Pu, M., Cai, Y., Lu, C., Li, Z., and Liang, G. (2021b). bHLH11 inhibits bHLH IVc proteins by recruiting the TOPLESS/TOPLESS-RELATED corepressors. Plant Physiol. doi: 10.1093/plphys/kiab540.

Long, T.A., Tsukagoshi, H., Busch, W., Lahner, B., Salt, D.E., and Benfey, P.N. (2010). The bHLH transcription factor POPEYE regulates response to iron defificiency in Arabidopsis roots. Plant Cell 22: 2219–2236.

Mori, S. (1999). Iron acquisition by plants. Curr Opin Plant Biol. 2: 250–253.

Pauwels, L., Barbero, G. F., Geerinck, J., Tilleman, S., Grunewald, W., Pérez, A. C., Chico, J. M., Bossche, R. V., Sewell, J., Gil, E., García-Casado, G., Witters, E., Inzé, D., Long, J. A., De Jaeger, G., Solano, R., and Goossens, A. (2010). NINJA connects the co-repressor TOPLESS to jasmonate signalling. Nature 464: 788–791.

Riaz, N., & Guerinot, M. L. (2021). All together now: regulation of the iron deficiency response. J Exp Bot. 72: 2045–2055.

Robinson, N. J., Procter, C. M., Connolly, E. L., and Guerinot, M. L. (1999). A ferric-chelate reductase for iron uptake from soils. Nature 397: 694–697.

Römheld, V., and Marschner, H. (1986). Evidence for a specific uptake system for iron phytosiderophores in roots of grasses. Plant Physiol. 80: 175–180.

Roberts, L. A., Pierson, A. J., Panaviene, Z., and Walker, E. L. (2004). Yellow stripe1. Expanded roles for the maize iron-phytosiderophore transporter. Plant Physiol. 135: 112–120.

Selote, D., Samira, R., Matthiadis, A., Gillikin, J. W., and Long, T. A. (2015). Iron-binding E3 ligase mediates iron response in plants by targeting basic helix-loop-helix transcription factors. Plant Physiol. 167: 273–286.

Sheftel, A. D., Mason, A. B., and Ponka, P. (2012). The long history of iron in the Universe and in health and disease. Biochim Biophys Acta. 1820: 161–187.

Szemenyei, H., Hannon, M., and Long, J. A. (2008). TOPLESS mediates auxin-dependent transcriptional repression during Arabidopsis embryogenesis. Science 319: 1384–1386.

Tanabe N, Noshi M, Mori D, Nozawa K, Tamoi M, Shigeoka S. (2019). The basic helix-loop-helix transcription factor, bHLH11 functions in the iron-uptake system in Arabidopsis thaliana. J Plant Res. 132: 93–105.

Tissot, N., Robe, K., Gao, F., Grant-Grant, S., Boucherez, J., Bellegarde, F., Maghiaoui, A., Marcelin, R., Izquierdo, E., Benhamed, M., Martin, A., Vignols, F., Roschzttardtz, H., Gaymard, F., Briat, J. F., and Dubos, C. (2019). Transcriptional integration of the responses to iron availability in Arabidopsis by the bHLH factor ILR3. New Phytol. 223: 1433–1446.

Trofimov, K., Ivanov, R., Eutebach, M., Acaroglu, B., Mohr, I., Bauer, P., and Brumbarova, T. (2019). Mobility and localization of the iron deficiency-induced transcription factor bHLH039 change in the presence of FIT. Plant direct 3: e00190.

Varotto, C., Maiwald, D., Pesaresi, P., Jahns, P., Salamini, F., and Leister, D. (2002). The metal ion transporter IRT1 is necessary for iron homeostasis and efficient photosynthesis in Arabidopsis thaliana. Plant J. 31: 589–599.

Vert, G., Grotz, N., Dédaldéchamp, F., Gaymard, F., Guerinot, M. L., Briat, J. F., and Curie, C. (2002). IRT1, an Arabidopsis transporter essential for iron uptake from the soil and for plant growth. Plant Cell 14: 1223–1233.

Wang, N., Cui, Y., Liu, Y., Fan, H., Du, J., Huang, Z., Yuan, Y., Wu, H., and Ling, H. Q. (2013). Requirement and functional redundancy of Ib subgroup bHLH proteins for iron deficiency responses and uptake in Arabidopsis thaliana. Mol Plant. 6: 503–513.

Yuan, Y., Wu, H., Wang, N., Li, J., Zhao, W., Du, J., Wang, D., and Ling, H. Q. (2008). FIT interacts with AtbHLH38 and AtbHLH39 in regulating iron uptake gene expression for iron homeostasis in Arabidopsis. Cell Res. 18: 385–397.

Yi, Y., and Guerinot, M. L. (1996). Genetic evidence that induction of root Fe (III) chelate reductase activity is necessary for iron uptake under iron deficiency. Plant J. 10: 835–844.

Yi, Y., Saleeba, J.A., and Guerinot, M.L. (1994). Fe uptake in Arabidopsis thaliana. Biochemistry of Metal Micronutrients in the Rhizosphere.

Zhang, J., Liu, B., Li, M., Feng, D., Jin, H., Wang, P., Liu, J., Xiong, F., Wang, J., and Wang, H. B. (2015). The bHLH transcription factor bHLH104 interacts with IAA-LEUCINE RESISTANT3 and modulates iron homeostasis in Arabidopsis. Plant Cell 27: 787–805.

