## Supplementaf information for "POPEYE directly regulates bHLH Ib genes and its own expression"

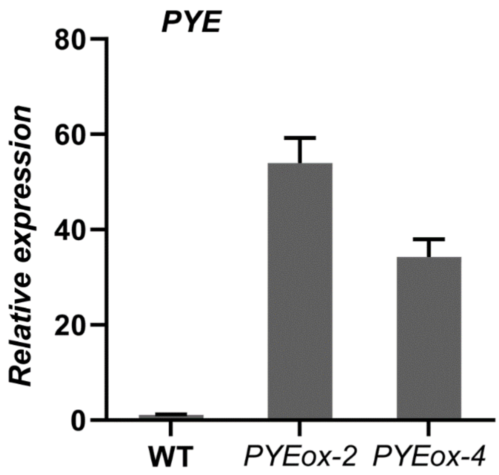


**Supplemental Figure S1.** Relative expression levels of *PYE* in *PYEox* plants. Four-day-old plants grown on +Fe were transferred to +Fe or -Fe medium for three days. Roots from 7-day-old seedling were harvested for the extraction of RNA and qRT-PCR.

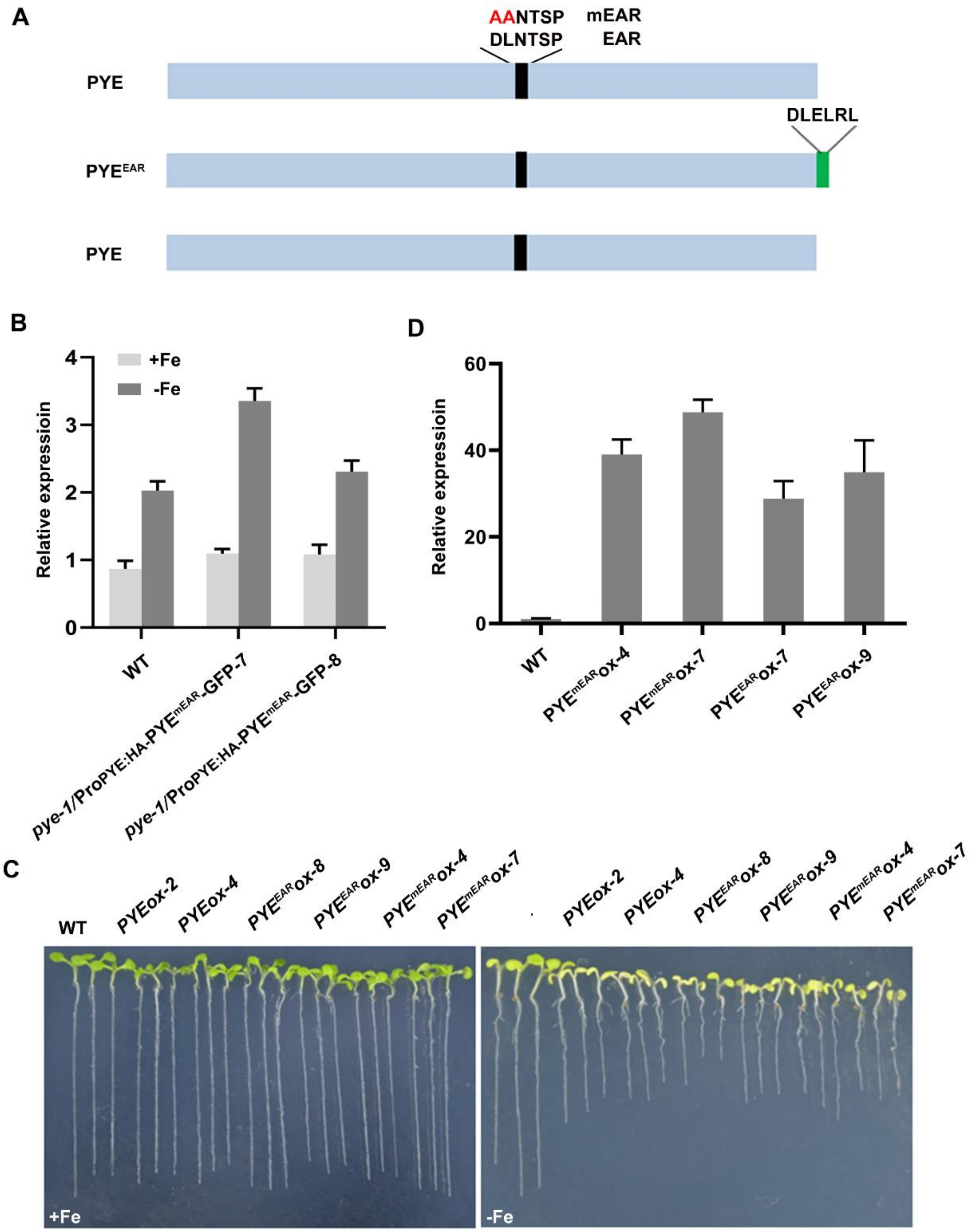


**Supplemental Figure S2.** The EAR of PYE is not required for its repression function.

(A) Schematic diagram of the various versions of PYE. The mutated amino acid is indicated in red. mEAR, the mutated EAR; $PYE^{EAR}$, the end of PYE was fused with an EAR domain.

(B) Relative expression levels of *PYE* in the $Pro_{PYE}$*:HA-*$PYE^{EAR}$*-GFP/pye-1* plants. Four-day-old plants grown on +Fe were transferred to +Fe or -Fe medium for three days. Roots were harvested for the extraction of RNA and qRT-PCR. The data represent means ± SD.

(C) Phenotypes of various transgenic plants. One-week-old seedlings grown on +Fe or -Fe medium.

(D) Relative expression levels of *PYE* in various transgenic plants. Four-day-old plants grown on +Fe were transferred to +Fe or -Fe medium for three days. Roots were harvested for the extraction of RNA and qRT-PCR. The data represent means ± SD.

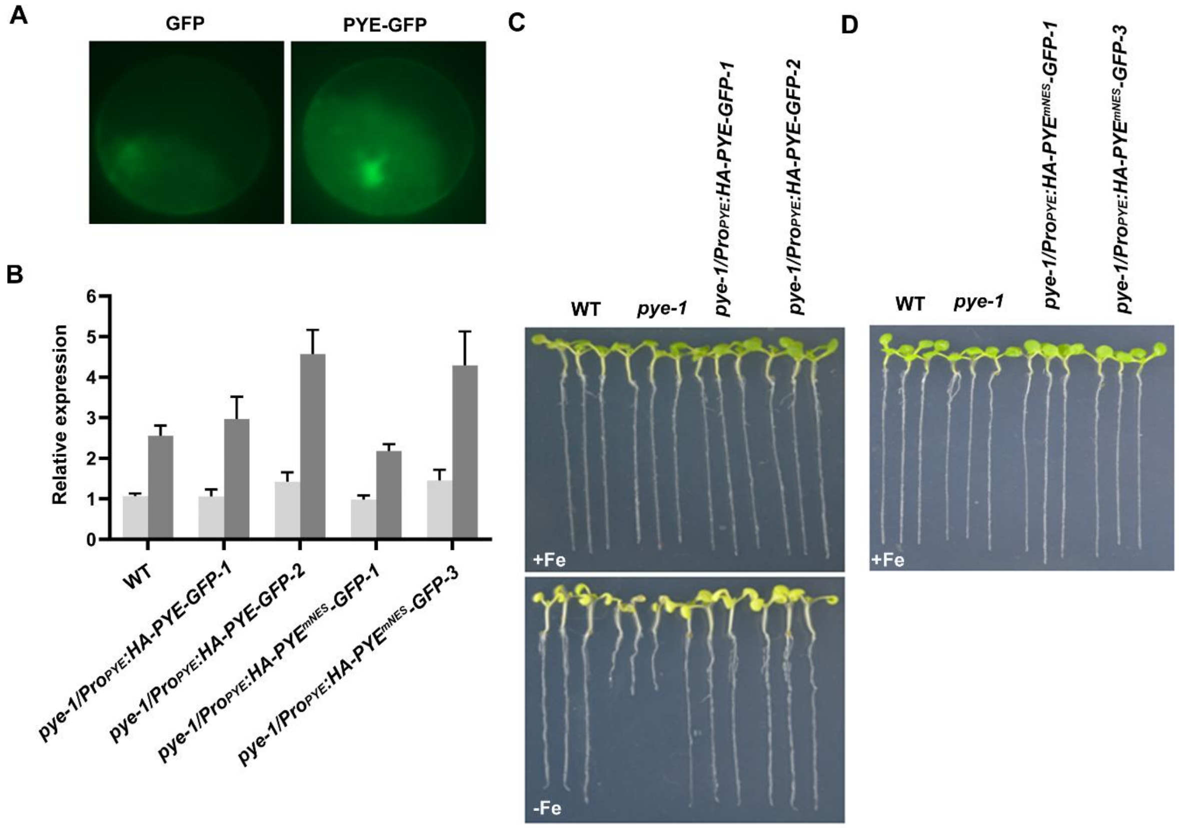


**Supplemental Figure S3.** The NES is required for PYE functions.

(A) Subcellular localization of PYE. GFP or PYE-GFP transformed into Arabidopsis mesophyll protoplasts. The GFP signal was visualized under a confocal microscope.

(B) Relative expression levels of *PYE* in various transgenic plants. Four-day-old plants grown on +Fe were transferred to +Fe or -Fe medium for three days. Roots were harvested for the extraction of RNA and qRT-PCR. The data represent means ± SD.

(C) Phenotypes of *pye-1/Pro*$_{PYE}$*:HA-PYE-GFP* plants. One-week-old seedlings grown on +Fe or -Fe medium.

(D) Phenotypes of *pye-1/Pro*$_{PYE}$*:HA-PYE*$^{mNES}$*-GFP* plants. One-week-old seedlings grown on +Fe medium.

**Supplemental Table S1**. Primers used in this study.

| | | | |
|---|---|---|---|
| PYE-F | ATAGGATCCATGGAGTACCCATACGACGTACCAGATTACGCTATGGTATCGAAAACTCCTTCTACAT | pOCA30 | *PYEox* |
| PYE-R | ATAGTCGACTCATTCACTGGCTTTCAGCC | | |
| $PYE^{mEAR}$-F | GTCGAAACCTGcCgcGAACACCTCTCCTGCACCCG | pOCA30 | $PYE^{mEAR}ox$ |
| $PYE^{mEAR}$-R | GAGGTGTTCgcGgCAGGTTTCGACTGATTCGCTCT | | |
| $PYE^{mNES}$-F | gcTGAATTAGCCGATACTCTTGAAgcGAATCAACAGAACAGTGGGAAAG | pOCA30 | $PYE^{mNES}ox$ |
| $PYE^{mNES}$-R | CgcTTCAAGAGTATCGGCTAATTCAgcGAAAAGCTCATTCAAATGCTC | | |
| PYE-F | ATAGGATCCATGGAGTACCCATACGACGTACCAGATTACGCTATGGTATCGAAAACTCCTTCTACAT | pOCA30 | $PYE^{mEAR}ox$ |
| $PYE^{EAR}$-R | ATAGTCGACTACAAACGGAGTTCGAGATCTTCACTGGCTTTCAGCCGCT | | |
| pPYE-PYE-F | GAAAGAATTCGAGCTCGCCCGGGCGAACCGCAAAACTATATATAGTA | p28-GFP | $Pro_{PYE}$:*HA-PYE-GFP* |
| pPYE-PYE-R | CTGGTACGTCGTATGGGTACTCCATCTTTGCTTTTATTACAGAACAAGA | p28-GFP | $Pro_{PYE}$:*HA-PY*$E^{mEAR}$*-GFP* |
| PYE-HA-F | ATGGAGTACCCATACGACGTACCAGATTACGCTATGGTATCGAAAACTCCTTCTACAT | p28-GFP | $Pro_{PYE}$:*HA-PY*$E^{mNES}$*-GFP* |
| PYE-GFP-R | GCCCTTGCTCACCATGGTTCTAGAGTCACTGGCTTTCAGCCGCTC | p28-GFP | |
| pPYE-F | TTgtcgaCGAACCGCAAAACTATATATAGTA | p28-GUS | $Pro_{PYE}$:*GUS* |
| pPYE-R | AAggatcCTTTGCTTTTATTACAGAACAAGA | | |
| GAD-PYE-F | TTTGAATTCATGGTATCGAAAACTCCTTCTACA | pGAD-T7 | *pGAD-PYE* |
| GAD-PYE-R | TTTGGATCCTCATTCACTGGCTTTCAGCC | | |
